# Biologically-informed machine learning identifies a new clinically-actionable bladder cancer subgroup characterised by NRF2 overactivity

**DOI:** 10.1101/2025.06.03.657659

**Authors:** Vlad V. Ungureanu, Richard T. Gawne, Simon C. Baker, Pontus Eriksson, David M. Halliday, Stephen L. Smith, Jennifer Southgate, Andrew S. Mason

## Abstract

Muscle-invasive bladder cancer is a diverse disease where subtyping is ambiguous. Gene expression profiling followed by unsupervised machine learning (ML) has broadened our understanding of tumour biology, but has failed to provide high-confidence clinically-actionable subgroups. To focus on tissue-specific urothelial biology, we generated co-expression networks from histologically normal bladder, including multiple differentiation states and prioritising transcription factors (TFs). This strategy revealed an emergent set of 98 TFs which we used to stratify The Cancer Genome Atlas bladder cancer cohort, revealing a subdivision of basal tumours characterised by the detoxification and glutaminolysis activity of NRF2, rendering them resistant to standard bladder cancer interventions. These 20 tumours (4.9%) expressed squamous markers, were highly aggressive (15% 2-year survival), and had signatures of active PI3K, MTOR and retinoic acid signalling. Intriguingly, only half of the subgroup had activating mutations in the NRF2/KEAP1 pathway, whilst half of putative driver *NFE2L2* mutations were excluded. This highlighted the importance of expression-based classification, particularly as re-analysis of *NFE2L2*-mutated lung cancer trial data showed only mutations consistent with our classification strategy responded to NRF2 inhibition. Our approach provides the first direct evidence that unsupervised ML can be biologically-informative in identifying clinically-actionable subgroups.

## Advancing practice: what does this study add?

This study provides a novel method using machine learning to find clinically-actionable groups of bladder cancer patients. We have moved beyond a data-only approach to prioritise gene sets with known relevance in healthy urothelium, finding a novel group of highly aggressive muscle invasive cancers which are eligible for treatment with NRF2 inhibitors. We propose that rather than attempting global classification schemes, the field should move to hypothesis-driven identification of bladder cancer subgroups optimised to confidently assign patients to a single personalised treatment regime..

## Patient summary

We used our understanding of healthy bladders to develop a new computational approach for finding subgroups of bladder cancer. This led us to discover a new group of patients who don’t respond to current bladder treatments, but would be eligible for a treatment already used in lung cancer. We can now use our new strategy to find more options for personalised treatment.

## Take-home message

We have demonstrated that integrating tissue-specific urothelial knowledge is essential for identifying clinically-actionable bladder cancer subgroups. Following this innovative strategy, we found a novel, highly-aggressive subgroup eligible for treatment with NRF2 inhibitors.

## Manuscript

Muscle invasive bladder cancer (MIBC) has high tumour mutational burden and the greatest diversity of driver mutations within The Cancer Genome Atlas (TCGA) ^1^. 5-year survival is approximately 40%, with little improvement despite the adoption of refined surgical techniques, platinum chemotherapies, and immunotherapies. Gene expression profiling followed by unsupervised machine learning (ML) was used throughout the 2010s to derive molecular subtypes, culminating in a “consensus” classification in 2020 ^2^. However, these subgroups indiscriminately subdivide MIBC across the continuum of differentiation and immune status (Figure 1A). We need to improve our computational approach in order to find clear patient groups which are clinically actionable.

**Figure 1.**
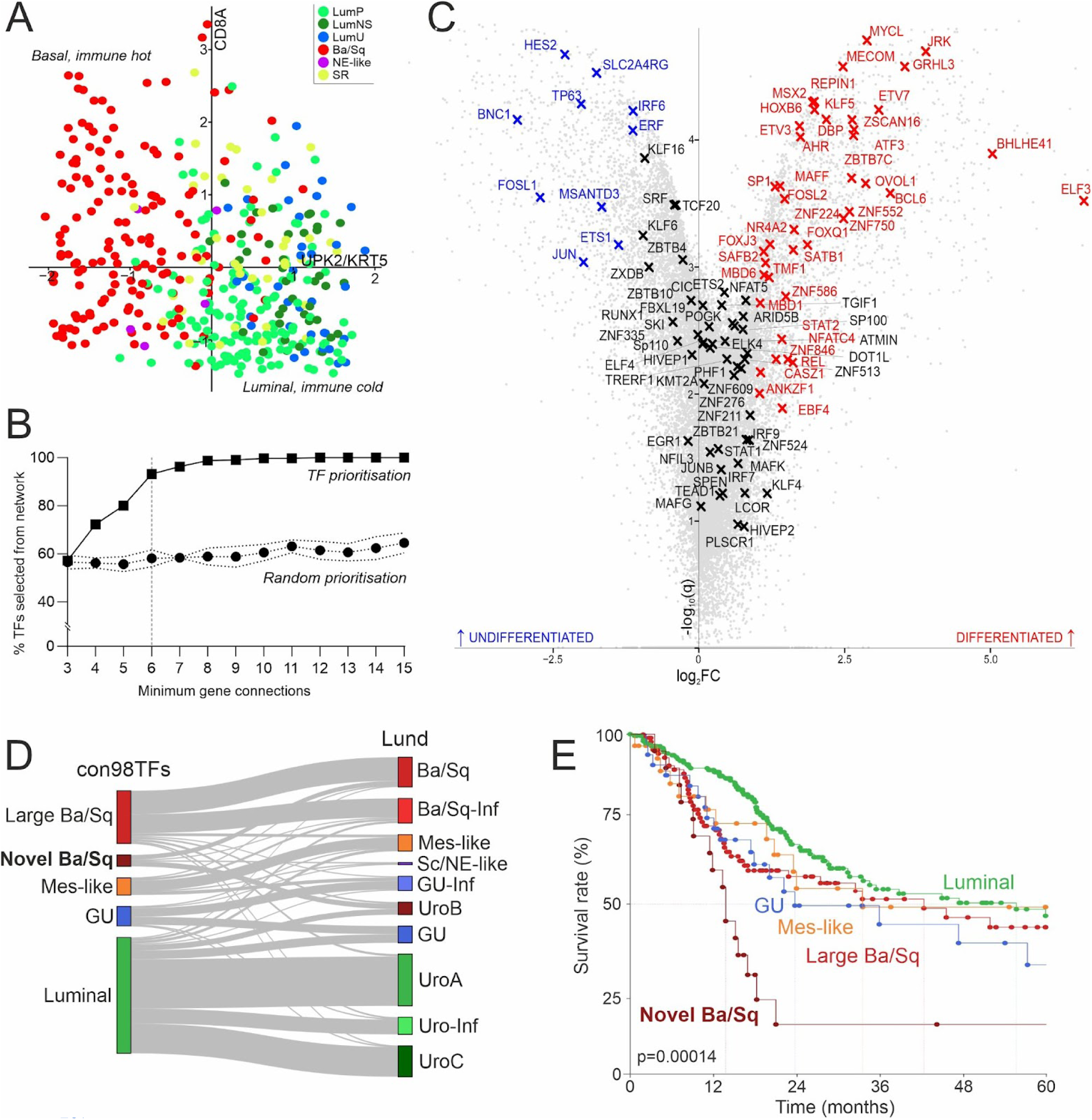
Network-based classification of TCGA MIBC cohort using histologically normal bladder reveals a novel Ba/Sq subdivision with very poor survival. A) MIBC molecular classification is largely based on differentiation and immune status, with z-scores of *UPK2*:*KRT5* ratio (x-axis) against *CD8A* z-score able to largely replicate the consensus classification ^2^. B) Increasing the minimum number of gene connections for transcription factors (TFs) increased the percentage of TFs within network community hubs eventually used for hierarchical clustering. This increase was not seen when other genes were randomly prioritised (repeated ten times with model standard deviation indicated). C) 98 TFs consistently emerged as highly connected hub genes, only 51.8% of which were associated with the differentiation status of normal urothelium. All labelled genes are in the list of conserved 98 TFs: red annotated genes upregulated in differentiated state; in blue those upregulated in undifferentiated cultures; and in black those not significantly associated with differentiation status. D) Hierarchical clustering of TCGA muscle invasive bladder cancer samples with these 98 TFs identified five major groupings largely consistent with the Lund classifier ^5^. E) The newly identified Novel Ba/Sq subgroup had very poor survival (15% 2-year survival).

Unsupervised ML typically uses bulk cancer gene expression data and selects the most highly-expressed and variable genes to separate samples into groups (Figure S1). Whilst conceptually unbiased, this approach includes impurity-based signatures from the wider tumour microenvironment, and fails to prioritise existing biological knowledge. To focus on tissue-specific urothelial biology, we constructed a gene co-expression network using RNA sequencing data derived from human urothelial cells isolated from histologically normal bladder (n=88; 47 independent donors). This dataset covered the major luminal/basal axis observed in MIBC, consisting of freshly isolated urothelium (n=23), de-differentiated and highly proliferative cultures akin to basal tumours (n=14), and re-differentiated biomimetic cultures (stratified, tight barrier-forming and quiescent; n=51) demonstrating luminal biology without environmental influences (Figure S2) ^3^. Unlike standard transcriptomic data preprocessing, we removed genes unless there was credible expression (TPM>1.5) in ≥90% of the samples, retaining genes which were integral for urothelial function regardless of differentiation state. We then selected the top 5000 ranked genes based on median-scaled variance, and constructed a co-expression network from Spearman correlation. Network complexity was vastly reduced by “selective edge pruning”: retaining only the best three correlations for each of the 5000 genes ^4^. We integrated biological understanding of gene regulation by varying the minimum number of connections allowed for transcription factors (TFs), selecting six based on the Elbow method (Figure 1B). TF prioritisation altered network topology, revealing highly-connected and biologically-consistent “hub genes” which underpinned communities within the network (Figures S3, S4).

Strikingly, 98 TFs consistently emerged as hub genes (Figure S4), and did not simply polarise based on differentiation status, validating our approach (Figure 1C). Hierarchical clustering of TCGA’s MIBC cohort (n=408) with these 98 TFs identified five subgroups highly consistent with the LundTax2023 classifier ^5^, with a Luminal supergroup (53.4%), Genomically Unstable (9.0%), Mes-like (7.9%) and a novel subdivision of Basal/Squamous (Ba/Sq) tumours (24.3% and 5.3%) independent of immune status (Figures 1D, S5, S6). Patients in the novel Ba/Sq group had a 2-year survival of just 15%, warranting further investigation (Figure 1E).

Differential expression and enrichment analysis of Novel Ba/Sq tumours revealed subgroup-specific detoxification activity driven by Nuclear factor erythroid 2-related factor 2 (NRF2; gene symbol *NFE2L2*), including glutathione conjugation and glutaminolysis (*GCLC*, *GCLM*, *GSTA1*, *SLC7A11*), *CYP4* and *UGT1* family members, and aldehyde and aldo-keto reductase activity (Figures 2A, 2B). Through such diverse detoxification strategies NRF2 confers resistance to reactive oxygen species generated by cellular stress, rendering these tumours resistant to chemotherapy and radiotherapy ^7^. Novel Ba/Sq tumours exhibited markers of squamous differentiation (consistent with histological metadata ^9^), resisting the luminal differentiation associated with the PI3K pathway activation (Figures 2B, 2C) and retinoic acid signalling ^7^. Tumour aggressiveness (Figure 1E) was consistent with immune exclusion (Figure S5; *IL6*, *CCL8*, *CCL13* downregulation in Figure 2A) and extracellular matrix dysregulation (*CEACAM5/6*, *CH13L2*, *L1CAM*). To validate the Novel Ba/Sq subgroup, a decision tree-based single sample classifier was developed on TCGA’s MIBC cohort (70:30 training:testing split with 96.58% training accuracy and 92.17% sensitivity; Figure S7) and used to classify the Lund s265 cohort ^5^. While rarer (2% of tumours, likely due to 56.7% of tumours being Ta/T1), the Novel Ba/Sq subgroup was identified in the validation cohort and exhibited a congruent transcriptomic profile (Figure S8).

**Figure 2.**
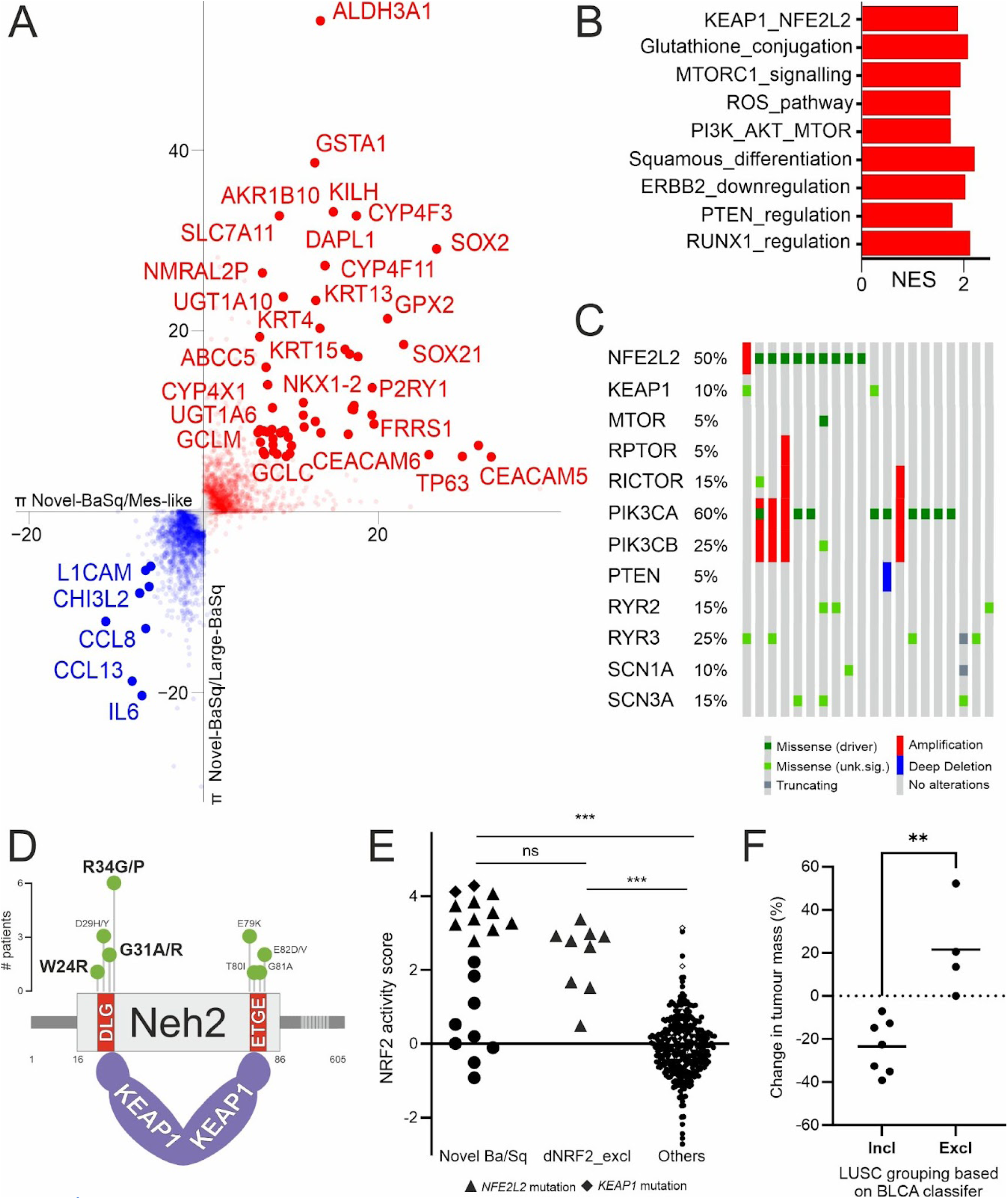
Novel Ba/Sq subgroup exhibits high NRF2, PI3K and retinoic acid activity. A) To determine the specific Novel Ba/Sq molecular profile we separately performed differential expression analysis between the Novel Ba/Sq and the Large Ba/Sq (y-axis) or Mes-like groups (x-axis). We generated π values for each gene in each condition, defined as the product of the log_2_FC and -log_10_ of the corrected significance value. Genes which scatter approximately to y=x are genes consistently up- (red) or downregulated (blue) in the Novel Ba/Sq subgroup. The Novel Ba/Sq-specific transcriptome was enriched for signatures of NRF2, PI3K and retinoic acid signalling. B) Broader upregulated pathways were highlighted by GSEA, including the clinically targetable MTOR and glutathione conjugation processes. C) Mutations (grouped by pathway) in the Novel Ba/Sq subgroup supported the overactivity of these pathways, including influencing intracellular calcium and sodium signalling. D) *NFE2L2* driver mutations are found in the Neh2-KEAP1 binding domain, with two binding sites of different affinities. Only tumours with *NFE2L2* driver mutations in the low affinity DLG were included in this Novel Ba/Sq group, but the literature-supported phenocopying mutations in the high-affinity ETGE binding site were excluded, as were D29 mutations which require mutated *KRAS* ^6,7^ not observed in any *NFE2L2*-mutated tumours. E) Detoxification and antioxidant activity was evident in the Novel Ba/Sq subgroup, but also in the tumours with driver *NFE2L2* mutations which were excluded from the Novel Ba/Sq group (dNRF2_excl). Mutation status alone was not sufficient to explain NRF2 activity. F) Stratification of *NFE2L2*-mutated non-small cell lung squamous carcinoma (LUSC) trial data ^8^ showed that only tumours with mutations consistent with the Novel Ba/Sq subgroup responded to treatment.

Canonically, NRF2 overactivity arises from ablating the direct inhibitory role of KEAP1. Mutations commonly alter the KEAP1 KELCH domains or the corresponding NRF2 Neh2 domain (Figure 2C). Any perturbation which disrupts NRF2/KEAP1 binding should enable NRF2 nuclear translocation and activation of antioxidant response elements ^10^. The Novel Ba/Sq subgroup contains half of the putative *NFE2L2* driver mutations in TCGA’s MIBC cohort, but all mutations in the KEAP1-high-affinity ETGE ‘hinge’ domain ^7^ were excluded (Figures 2C, 2D; also excluded in the validation cohort). Differential expression analysis revealed that these excluded tumours had no upregulation of glutaminolysis, MTOR signalling or the high retinoic acid activity (TCGA RARɣ regulon NES=1.70 q=10^-10^) seen in the Novel Ba/Sq subgroup, although squamous changes and ROS pathway activity were present across all NRF2 mutated tumours (Figure S9). Full NRF2 pathway activation is more broadly observed, suggesting context-dependent signalling beyond what mutation status would predict, and requires expression-based identification (Figures 2C-E, S7, S8, S10).

Encouragingly for patients with this aggressive form of MIBC, NRF2 pathway inhibitors have been developed and used in trials for refractory non-small cell lung squamous cell carcinoma (LUSC), supporting an expedited route to the clinic for NRF2 overactivity in MIBC. While trial success has been mixed, re-analysis of outcomes from the phase II NCT02417701 LUSC trial ^8^ showed that tumours with NRF2 mutations consistent with our Novel Ba/Sq subgroup all responded to treatment (median 22.6% reduction in tumour mass), whereas tumours with NRF2 hinge domain mutations either did not respond or progressed (Figure 2F). This means that our NRF2 activity-based identification strategy has broader relevance for improving patient inclusion and treatment efficacy in LUSC and other cancers with overactive NRF2.

This study demonstrates that we can use a network representation of gene regulation in normal urothelium to biologically-inform unsupervised ML strategies and identify a clinically-actionable bladder cancer subgroup. Whilst an all-encompassing single MIBC classification is appealing, we conclude that the field should move to hypothesis-driven subgroup identification, where we can confidently recommend more personalised treatment.

## Author contributions

Conception and design: Mason

Acquisition of data: Mason, Ungureanu, Eriksson

Analysis and interpretation of data: Ungureanu, Mason, Gawne, Eriksson, Baker, Southgate

Drafting of the manuscript: Ungureanu, Mason

Critical revision of the manuscript for important intellectual content: Ungureanu, Mason, Baker, Southgate

Statistical analysis: Ungureanu, Mason

Obtaining funding: Mason, Smith, Southgate

Administrative, technical, or material support: None

Supervision: Mason, Southgate, Halliday, Smith

Other: None

## CRediT authorship contribution statement

Vlad V. Ungureanu: Data curation, Formal analysis, Investigation, Methodology, Software, Visualization, Writing – original draft

Richard T. Gawne: Data curation, Formal analysis

Simon C. Baker: Formal analysis, Investigation, Validation, Writing – review & editing

Pontus Eriksson: Data curation, Formal analysis, Investigation

David M. Halliday: Project administration, Supervision, Writing – review & editing

Stephen L. Smith: Funding acquisition, Project administration, Supervision, Writing – review & editing

Jennifer Southgate: Funding acquisition, Project administration, Resources, Supervision, Writing – review & editing

Andrew S. Mason: Conceptualization, Data curation, Formal analysis, Funding acquisition, Investigation, Project administration, Resources, Software, Supervision, Validation, Visualization, Writing – original draft, Writing – review & editing

## Ethical approval of studies

All gene expression and mutation data were reanalysed from public repositories, with methodology and scope approved by The University of York Biology Ethics Committee (AM202410). Authorised Access to raw RNA sequencing data within The Cancer Genome Atlas was granted through dbGap project #25297 and processed on The University of York’s High Performance Computing Cluster ‘Viking’ in accordance with local and eRA data management policies.

## Data availability

All raw RNA sequencing data analysed in this study is publicly available in FASTQ format. The healthy urothelial RNA sequencing data is available directly from SRA projects PRJNA1265789 and PRJNA1266101. Authorised Access data within The Cancer Genome Atlas is available by application through dbGaP (phs000178). Gene-level expression data from the validation Lund s265 cohort is available on Zenodo (available from: https://zenodo.org/records/10362517). Mutation and survival data for TCGA is available through cBioPortal (available from: https://www.cbioportal.org/study/summary?id=blca_tcga_gdc).

All code used for the analysis of these datasets is available on Github: https://github.com/asmasonomics/biology-driven-uML

## Acknowledgements

We would like to acknowledge the University of York high performance compute facility “Viking” (compute resource) as well as the supportive interdisciplinary environment of the Jack Birch Unit (core funded by York Against Cancer) and York Biomedical Research Institute. We also thank Dr Matthew Care, author of PGCNA, for many helpful discussions throughout Vlad V. Ungureanu’s PhD.

## Supplementary methods

### Transcriptomic profiling of histologically normal bladder urothelium

mRNA sequencing data (n=88) previously generated by our unit from histologically normal urothelium was publicly available under accessions PRJNA1265789 and PRJNA1266101. Briefly, bladder biopsies had been collected from patients undergoing rigid or flexible cystoscopy for non-malignant bladder disease with lower urinary tract symptoms. Each dataset was derived from trypsin-isolated, high-purity urothelial cell sheets in one of three differentiation states, as described previously ^3^: freshly isolated “*in situ*” (n=23); actively proliferating “undifferentiated” (n=14); and biomimetic, barrier-forming and quiescent “differentiated” (n=51). The dataset includes a total of 47 independent donors, with some donors represented in two (n=11) or all three (n=11) differentiation states. All datasets were Illumina paired-end polyA libraries, using either the HiSeq4000 (100bp PE) or NovaSeq6000 (150bp PE) instruments.

Raw sequencing data were pseudoaligned to the human Gencode v42 comprehensive transcriptome annotation using kallisto v0.46.1 ^11^ and summarised to gene-level transcripts per million (TPM) with tximport v1.24.0 ^12^. Short non-coding (i.e. technical noise) and mitochondrial genes were removed from the dataframe and TPMs recalculated to total 10^6^. To retain genes which were informative across all differentiation states, an aggressive filtering strategy was employed, requiring a gene to be expressed with a TPM > 1.5 in 90% of the samples. Unlike standard machine learning pipelines, this removed genes with high variance if they were not expressed in the majority of samples. The remaining 14027 features were ranked by highest relative variance (cohort standard deviation divided by median value), and then the top 5000 genes were selected for network construction.

### Derivation of gene co-expression network from histologically normal bladder urothelium

Co-expression networks were constructed using our own network library, iNet, inspired by PGCNA2 ^4^. First, the selected top 5000 most informative genes were log_2_(TPM+1) transformed and used to construct an all-vs-all Spearman correlation matrix generating a network with 1.25×10^7^ edges. “Selective edge pruning” was employed to initially retain the best three non-self correlations (edges) from each gene (node), retaining network information but hugely reducing complexity ^4^. Importantly, genes can have more than three edges within the resultant network (so-called “hub nodes”) as any one gene may be present in the top three correlations of many different genes. Community detection and gene selection has been significantly improved through our iNet library with two specific features. First, we employed degree corrected stochastic block model (dcSBM) ^13^ rather than the often-employed Louvain or Leiden community detection tools, as these have been shown to derive communities from modelled noise, and are unable to identify small communities, particularly as network complexity increases ^13,14^. Second, we simplified the Module Connectivity “ModCon” score used to select the 100 most informative genes from each identified network community, to reflect the use of only a single dataset during network construction: using median expression for each gene within the dataset.

### Implementing biological prioritisation within the network

To prioritise hypothesised sets of biologically significant genes into the co-expression network, we modified the minimum and maximum permissible edge degree during selective edge pruning. To demonstrate this strategy we first prioritised the 1639 genes classified as transcription factors (TFs) by The Human Transcription Factors database v1.01 ^15^, of which 325 were present within the 5000 most informative genes. Minimum degree was varied in 1-degree increments between 3 (default) and 15, identifying a plateau after a degree of 10, with the majority of model gains by a minimum degree of 6. To understand the explanatory effects of prioritising TFs, this model was compared with prioritising an equal number of randomly selected genes, following the same 1-degree increments, repeated ten times.

The association of any TF with a particular differentiation state within the histologically normal urothelial dataset was assessed using the full TPM data, and by performing differential expression using donor sets with matched biomimetic/differentiated and proliferative/undifferentiated states (n=3).

### Remapping TCGA MIBC transcriptomic data

Aligned RNA sequencing data for 408 primary bladder cancer samples from The Cancer Genome Atlas were accessed under dbGaP project 25297. In the four cases with multiple sequencing runs, the rerun alignment files were selected as the representative sample. Data were downloaded using the Genomic Data Commons command line client v1.5.0 (available from: https://gdc.cancer.gov/access-data/gdc-data-transfer-tool). FASTQ files were regenerated from sample BAM files using samtools v1.10 ^16^ to exclude secondary and supplementary alignments, followed by BEDTools v2.27.1 bamToFastq ^17^. No additional quality control steps were performed on the extracted read files, and reads were pseudoaligned and quantified to TPMs as described above.

### Stratification of TCGA MIBC samples using transcription factors identified from network analysis of healthy urothelium

Gene-level TPMs from TCGA’s MIBC cohort were log_2_(TPM+1) transformed and quantile normalised before extracting data solely from the 98 TFs identified from the co-expression network analysis described above.These data were used for agglomerative hierarchical clustering with average linkage and the 1-Pearson correlation distance. Derived tumour sample clusters representing less than 1% of the total cohort size (i.e. four samples or fewer) were removed, resulting in a final dataset of 378 samples distributed across five groups.

Differentially-expressed genes between all pairwise comparisons of the five groups were identified using the Sleuth v0.30.0 ^18^ implementation of the likelihood ratio test, generating Benjamini-Hochberg corrected significance values. π-values ^19^ were derived for all genes within each comparison from the product of the TPM-derived log_2_ fold change values and the -log_10_ of the corrected significance values, where corrected significance values had a maximum allowable value of 0.99 to reduce the number of π-values which equalled zero. Individual comparison π-values were used to rank genes for pre-ranked gene set enrichment analysis using GSEApy v0.10.2 (available from: https://github.com/zqfang/GSEApy) against the Molecular Signatures Database (MSigDB) ^20–22^ and Reactome collections ^23^. Comparison-specific gene ranking was performed by determining a reference point with coordinates based on the largest π-values in the included pair of relevant comparisons, then measuring the shortest linear distance between each gene and the reference point, providing a rank order.

### Expression dataset annotation with existing bladder molecular classifiers

Cancer and histologically normal gene expression data was used to classify all samples according to the consensus (available from: https://github.com/cit-bioinfo/consensusMIBC ^2^) and LundTax2023 (available from: https://github.com/LundBladderCancerGroup/LundTaxonomy2023Classifier ^5^) molecular classifiers in R v4.3.3 (R Core Team, 2024). Samples were also compared with the original cohort classifications^9^.

### Correlation with exome-called mutation data

Sample-matched exome data for TCGA’s MIBC cohort (TCGA, GDC) were accessed via the cBioPortal API using cbioportalR ^24^ using Maftools v3.18 ^25^ and the OncoKB annotations ^26^.

### Development of a single sample classifier for the derived subgroups

The final TCGA MIBC gene expression dataset of 378 samples stratified in the hierarchical clustering was split into a training and test dataset at a ratio of 70:30, maintaining the proportions of each of the five identified subgroups. The random forest decision tree underlying the classifier was constructed using the multiclassPairs v0.4.3 ^27^ package in R to train a top-score pairs algorithm with a “one vs one” strategy implementation of Dunn’s statistical test to find the pairwise gene rules specific to each identified subgroup. The model was trained on the top 10000 most differentially expressed genes, allowing non-variant “pivot” genes to be included within rules.

### Implementation of the single sample classifier on the Lund s265 bladder cancer cohort

RNA sequencing data from the Lund s265 bladder cancer cohort ^5^ was used for classifier and subgroup biology validation. Individual sample abundance files (kallisto v0.48.0 pseudoaligned to the Gencode v39 annotation) were downloaded from https://zenodo.org/records/10362517 and summarised to protein coding gene-level TPMs using code provided within the Zenodo repository. Three samples were removed due to relatively poor sequencing depth, and one non-urothelial primary. Unlike TCGA, the remaining 261 samples represented diverse disease stages: 16 Ta, 132 T1, 112 T2+, 1 Tx. Derived TPMs were classified using the single sample classifier described above. Subgroup differential expression and gene set enrichment analysis were performed as described above.

### Mutation calling from the Lund s265 RNA sequencing data

RNA sequencing reads were aligned to the GRCh38.p14 primary human assembly using STAR v2.7.10a ^28^ through the NFcore ^29^ RNAseq pipeline v3.14 (available from: 10.5281/zenodo.1400710) with Ensembl v110 ^30^ gene annotations. Single nucleotide variants were called using GATK v4.4.0.0 HaplotypeCaller ^31^ and annotated with SNPEff v5.2f27 ^32^ and VEP ^33^. Non-synonymous variants called within the canonical *NFE2L2*, *KEAP1* and *PIK3CA* transcripts were manually inspected to remove likely germline contamination and extract relevant hotspot and driver mutations.

## Supplementary figures

**Figure S1.**
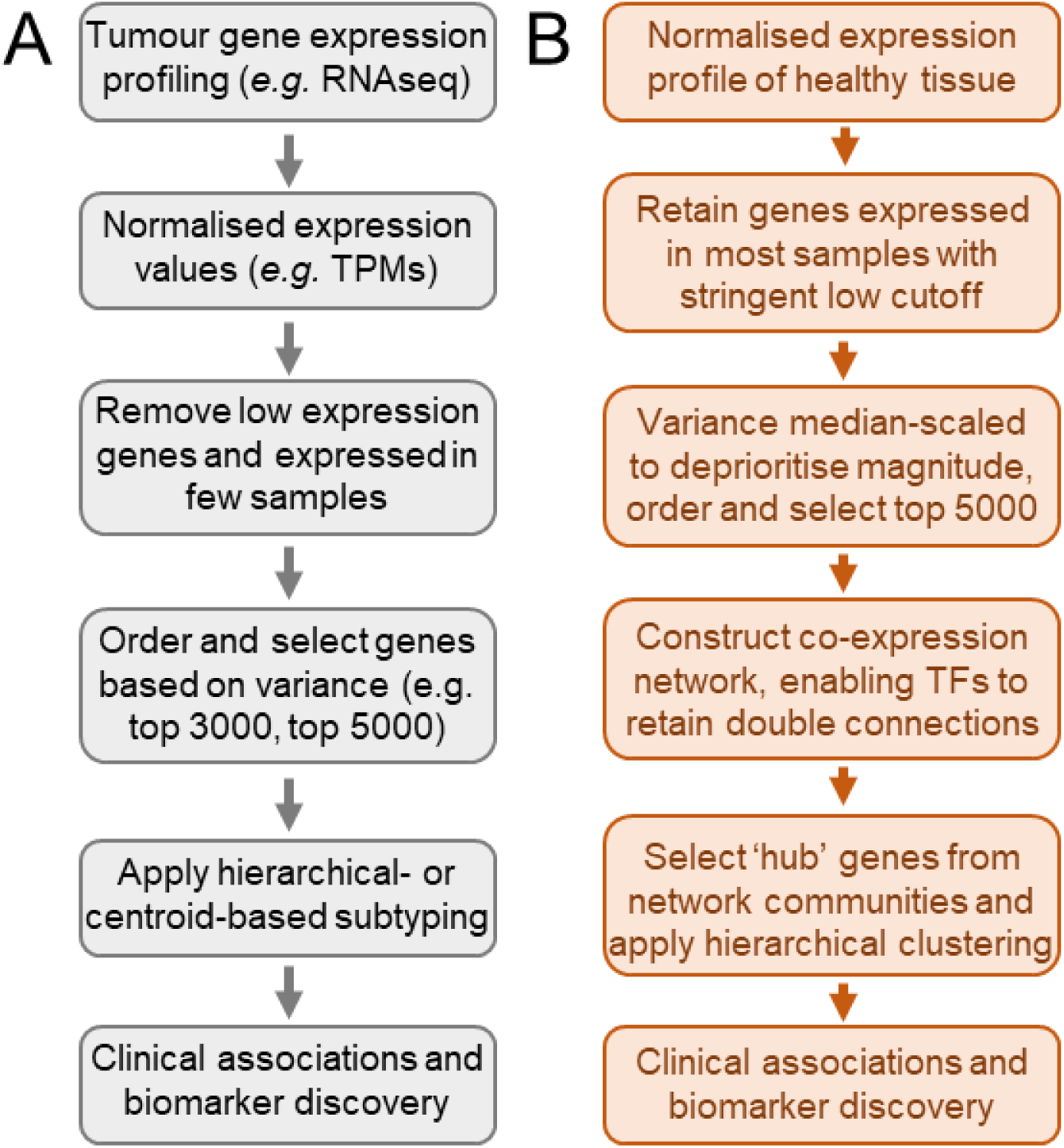
Strategies for sample stratification using unsupervised machine learning with gene expression data. A) Example simplified workflow for processing tumour gene expression data (typically now RNA sequencing, previously microarray). Gene selection biased towards genes which separate out a few samples and/or have high magnitude (generating high variance in both cases). Tumour biopsies often do not contain only cancer cells leading to the presence of cells from the wider tumour and/or paracancerous tissue within the transcriptome. These methods determine “informative” genes *de novo*. B) Summary workflow for our biologically-informed network representation strategy as described in the main manuscript.

**Figure S2.**
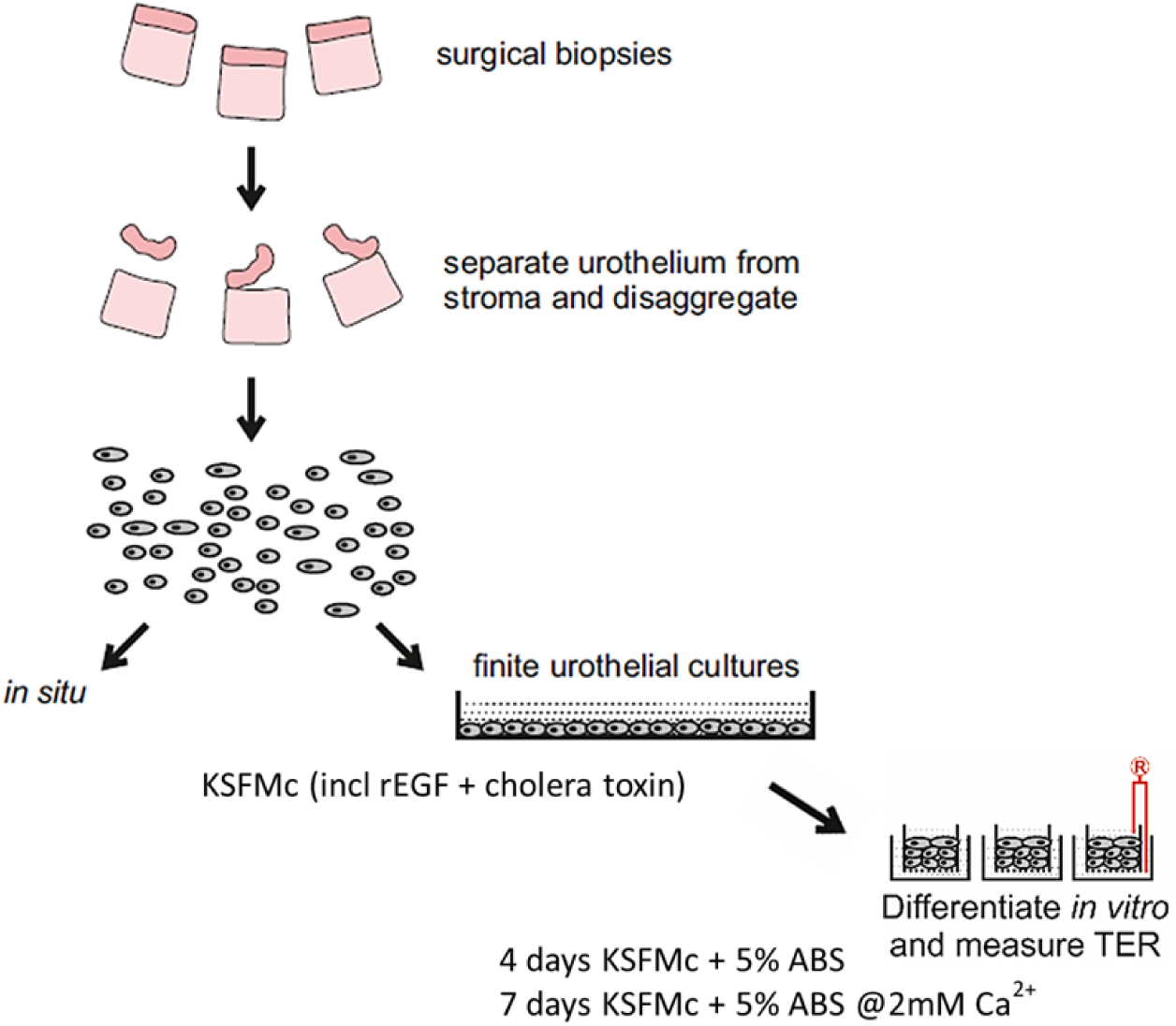
Normal Human Urothelium differentiation states and culture conditions. Histologically normal human urothelium sampled from surgical biopsies from patients with lower urinary tract symptoms was isolated from underlying stroma and disaggregated to a single suspension of almost pure urothelial cells. Samples to best reflect the patient’s urothelium (“*in situ*”) were processed for RNA immediately. Otherwise, urothelial cells would be cultured in keratinocyte serum free medium (KSFM) with recombinant epidermal growth factor (rEGF) and cholera toxin (together KSFM “complete”: KSFMc). After expansion of these non-transformed and finite cultures, cells can be redifferentiated by supplementing the media with adult bovine serum (ABS), seeding onto semi-permeable membranes and then increasing the concentration of Ca^2+^, as described previously^3^. Re-differentiated cultures form tight, physiological barriers in excess of 1000Ω.cm^2^, measured by transepithelial electrical resistance (TER).

**Figure S3.**
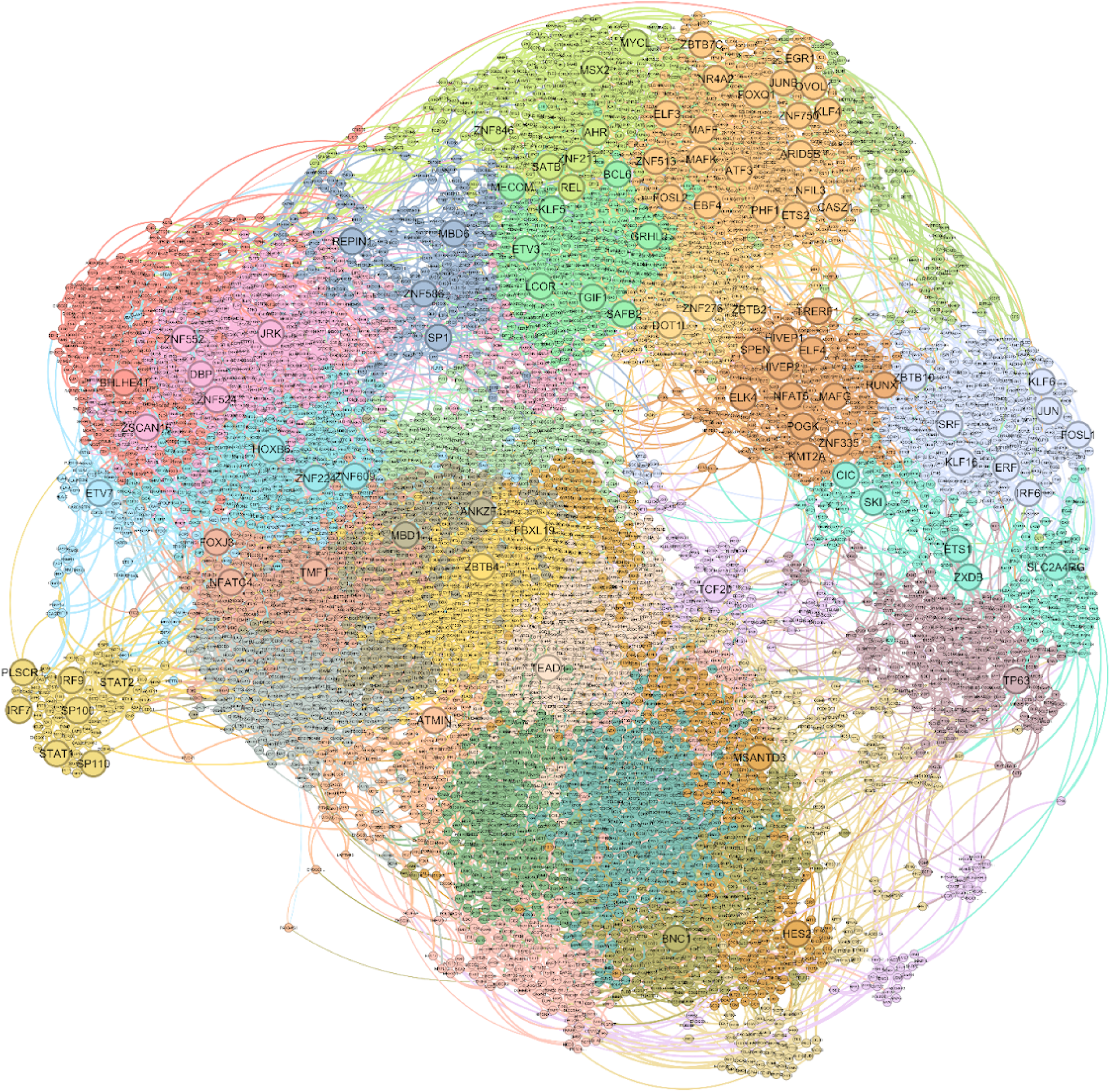
Representative urothelial network. Gene coexpression network where transcription factors were allowed to have at least six edges. Colours represent communities identified by degree corrected stochastic block model (dcSBM). Position of the emergent 98 TFs (Figures 1B, 1C, S4) indicated by larger circles.

**Figure S4.**
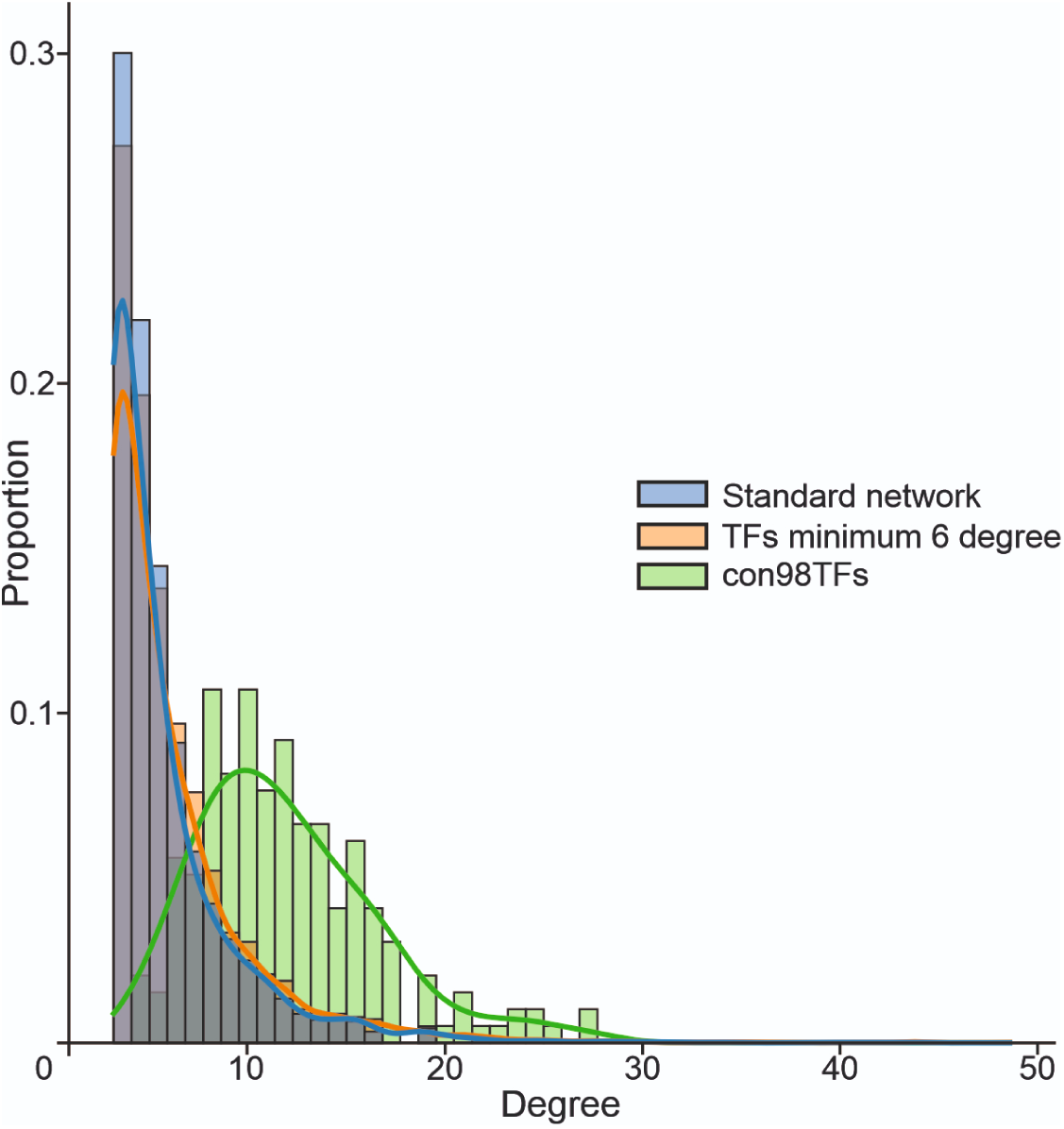
Degree distribution within standard and prioritised networks. Increasing the minimum degree of TFs to 6 significantly shifted network degree distribution (MWU 1.17x10^7^, p=2.50x10^-8^; KS p=1.10x10^-7^). The emergent TFs had inherently higher degree even without prioritisation (vs-standard MWU 1.05x10^5^, p=9.34x10^-81^; vs-min6 MWU 1.24x10^5^, p=1.42x10^-72^).

**Figure S5.**
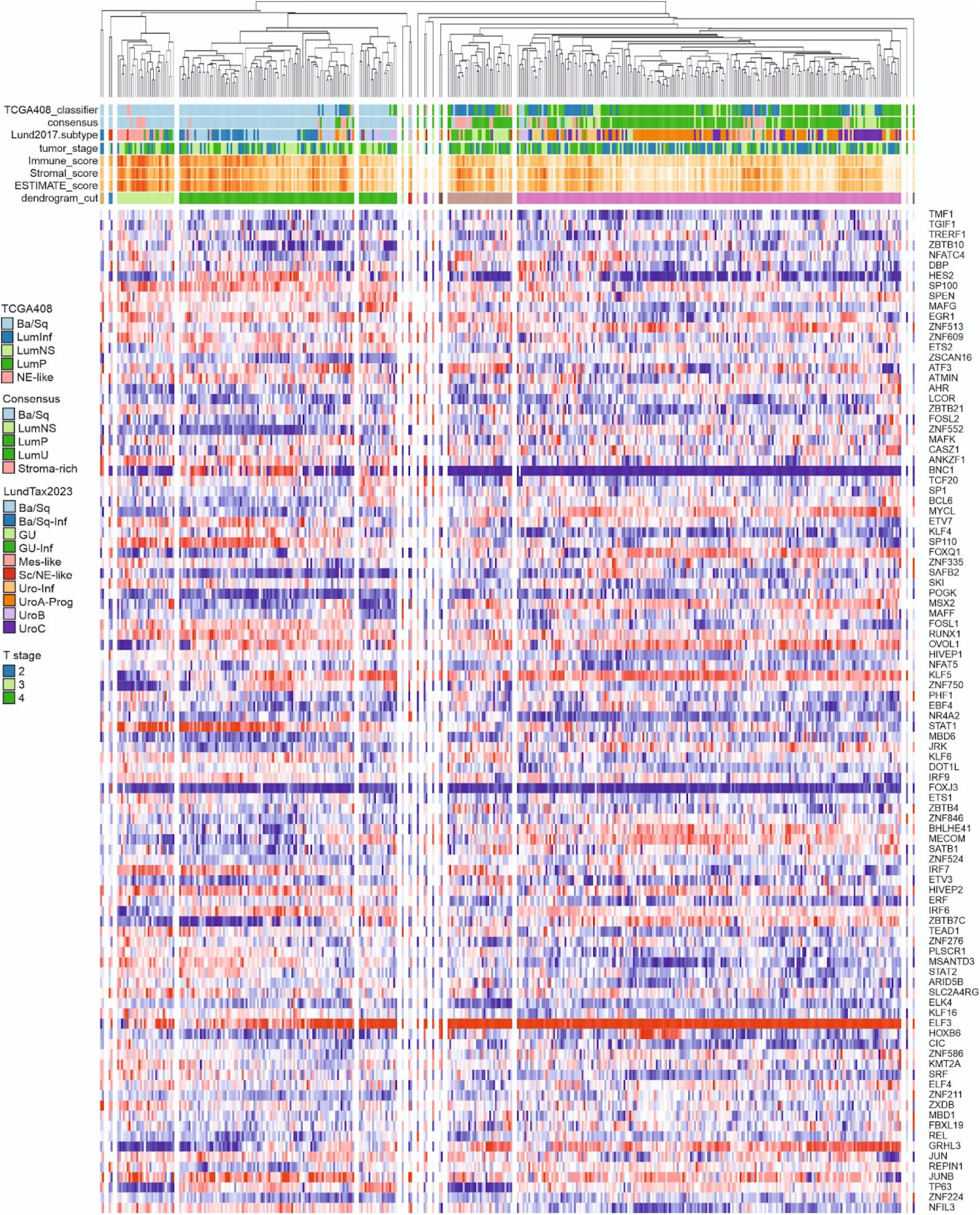
Hierarchical clustering of TCGA MIBC cohort using the emergent 98 transcription factors. TPMs for each gene were log_2_(TPM+1) transformed, normalised by the quantiles, and then used for agglomerative clustering using the 1-Pearson correlation distance with average linkage. The dendrogram cut was considered visually, based on major subdivisions whilst excluding clusters which represented less than 1% of the total cohort. Samples were further annotated with the molecular classifications from the 2017 TCGA paper^9^, MIBC consensus^2^, and LundTax2023^5^, as well as the Immune, Stromal and ESTIMATE scores included in TCGA MIBC cohort metadata^9^.

**Figure S6.**
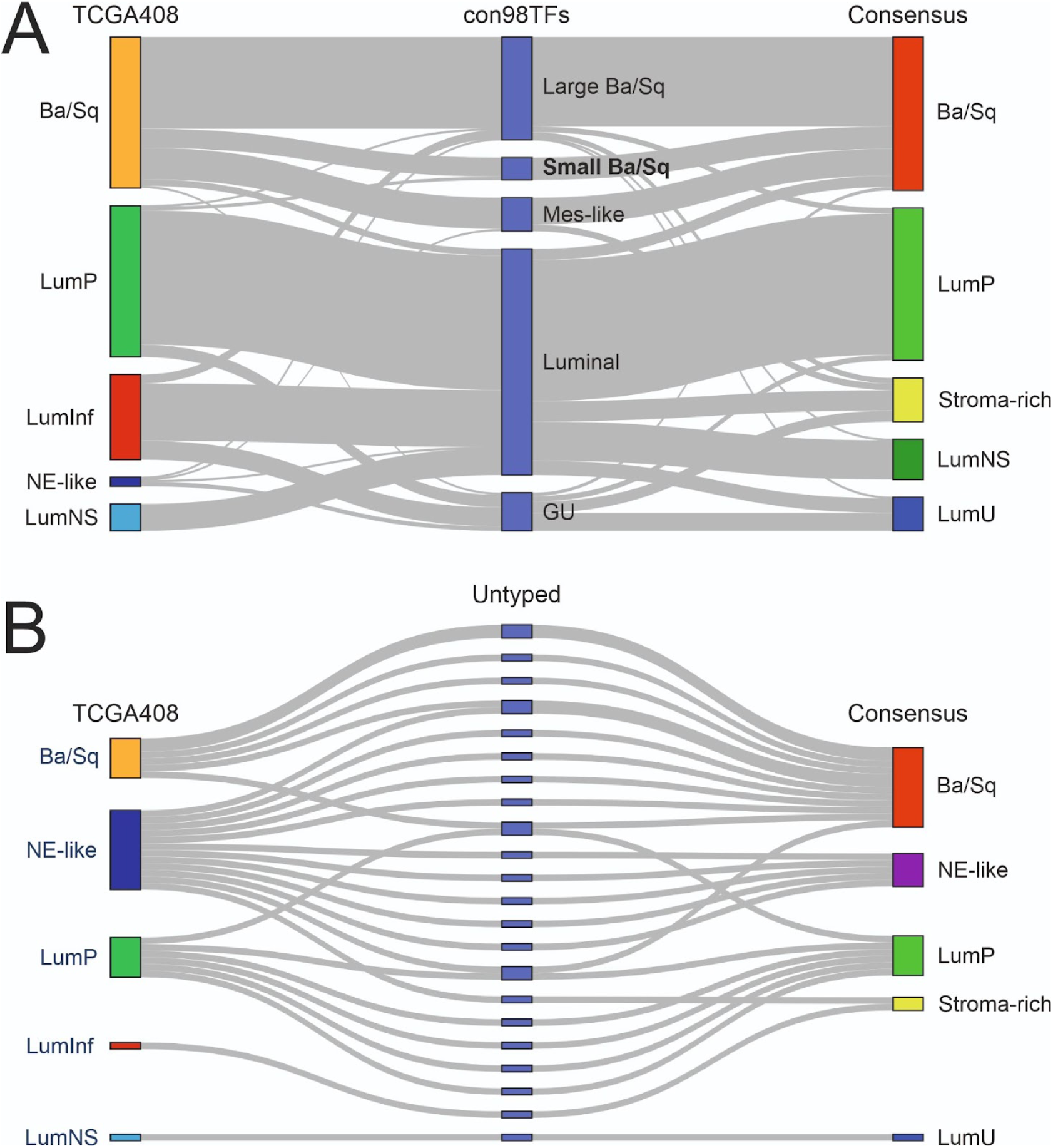
Comparison of sample classifications. A) Classification of 378 TCGA MIBC tumours following our strategy compared with TCGA’s original classification^9^ and the consensus classification^2^. Note that none of the NE-like tumours were classified in this strategy. B) Classification of samples which were removed as outliers as singlets or clusters with fewer than 4 samples in the hierarchical clustering (Figure S6).

**Figure S7.**
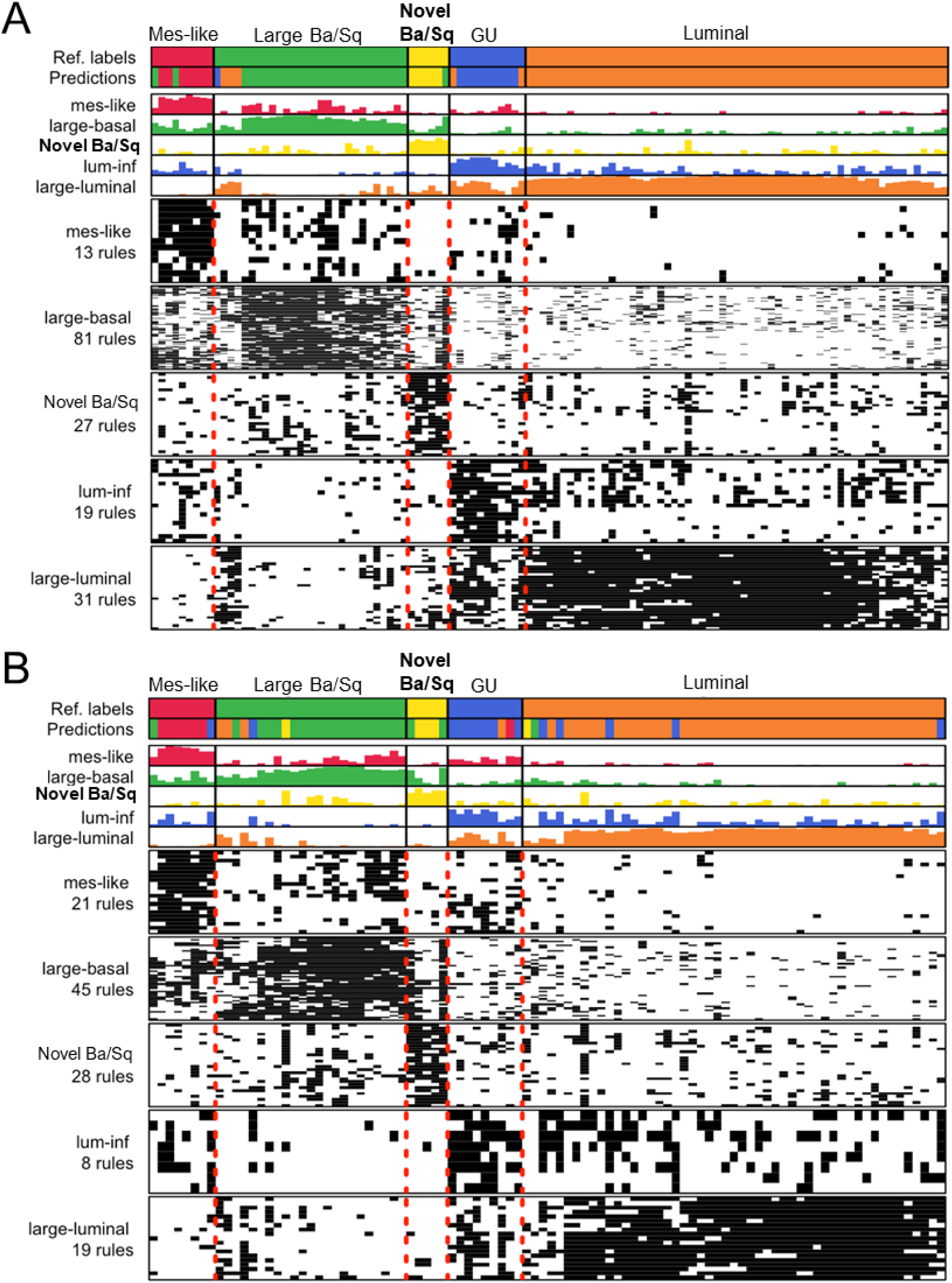
Development of Single Sample Classifier on TCGA MIBC cohort. Classifier was constructed using random forest decision tree based on top-score pairs algorithm from a “one vs one” implementation to find pairwise gene rules specific to each identified subgroup. Data were 70:30 split into training (A) and test (B) subsets with equal proportions of each identified subgroup. The model achieved 96.58% training accuracy and 92.17% sensitivity on the test data.

**Figure S8.**
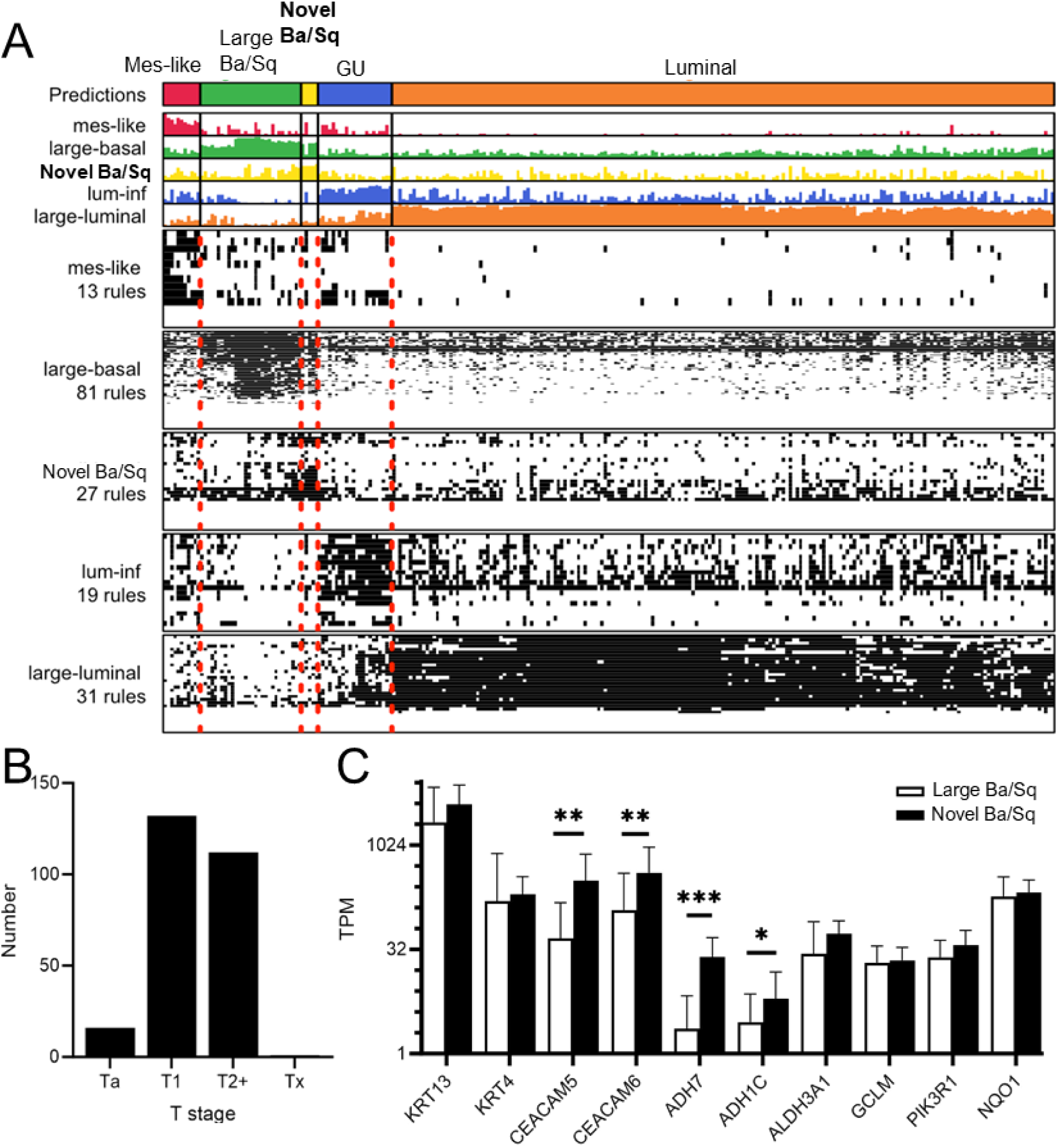
Lund s265 as a validation cohort for the novel Novel Ba/Sq subgroup. A) Our single sample classifier identified 5 samples consistent with the Novel Ba/Sq subgroup. As this appears likely a feature of more advanced disease, it is not unexpected to see a low proportion given the cohort T stage (B). C) Retinoic acid activity markers observed in TCGA were up in the Lund Novel Ba/Sq group, with reactive oxygen flat as expected. One sample was almost classified as Mes-like (A) and this was the most different to the other four, with TPM≈0 for *KRT13* and *KRT4* and low values for the *ADH* genes. *NFE2L2* mutations were only observed in three tumours and none in the Novel Ba/Sq, but this was a positive as mutations were either not drivers or fell in the ETGE domain excluded from our group.

**Figure S9.**
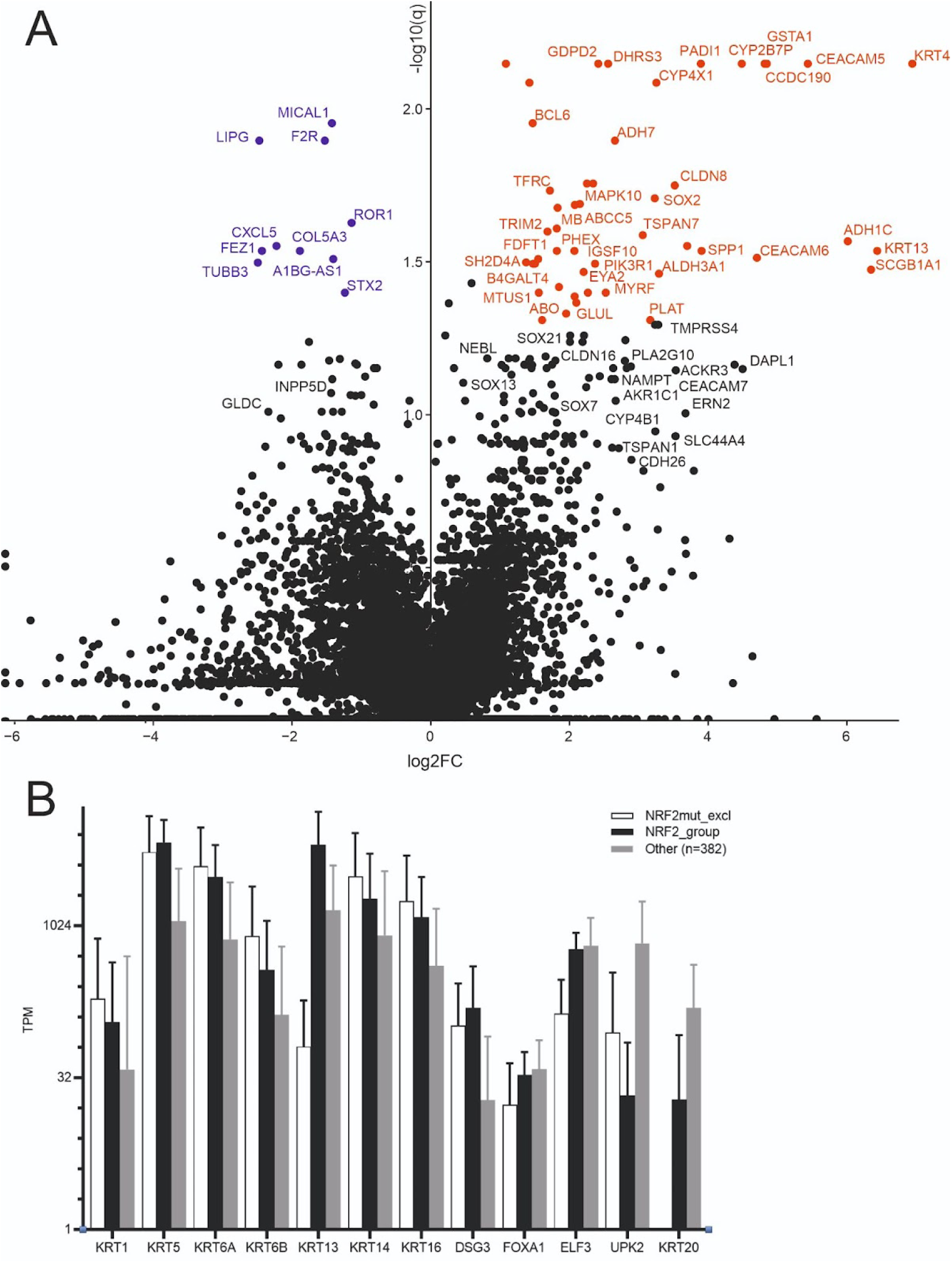
Transcriptomic differences between *NFE2L2* DLG and ETGE region mutated MIBC cancers. A) DLG region mutated tumours (n=10) had transcriptomic signatures consistent with elevated glutaminolysis, retinoic acid signalling and retinol metabolism, and greater invasiveness. B) DLG region mutated tumours and ETGE region mutated tumours (n=9) do not significantly differ for transitional or luminal markers, suggesting retinoic acid signalling was not promoting a more luminal phenotype.

**Figure S10.**
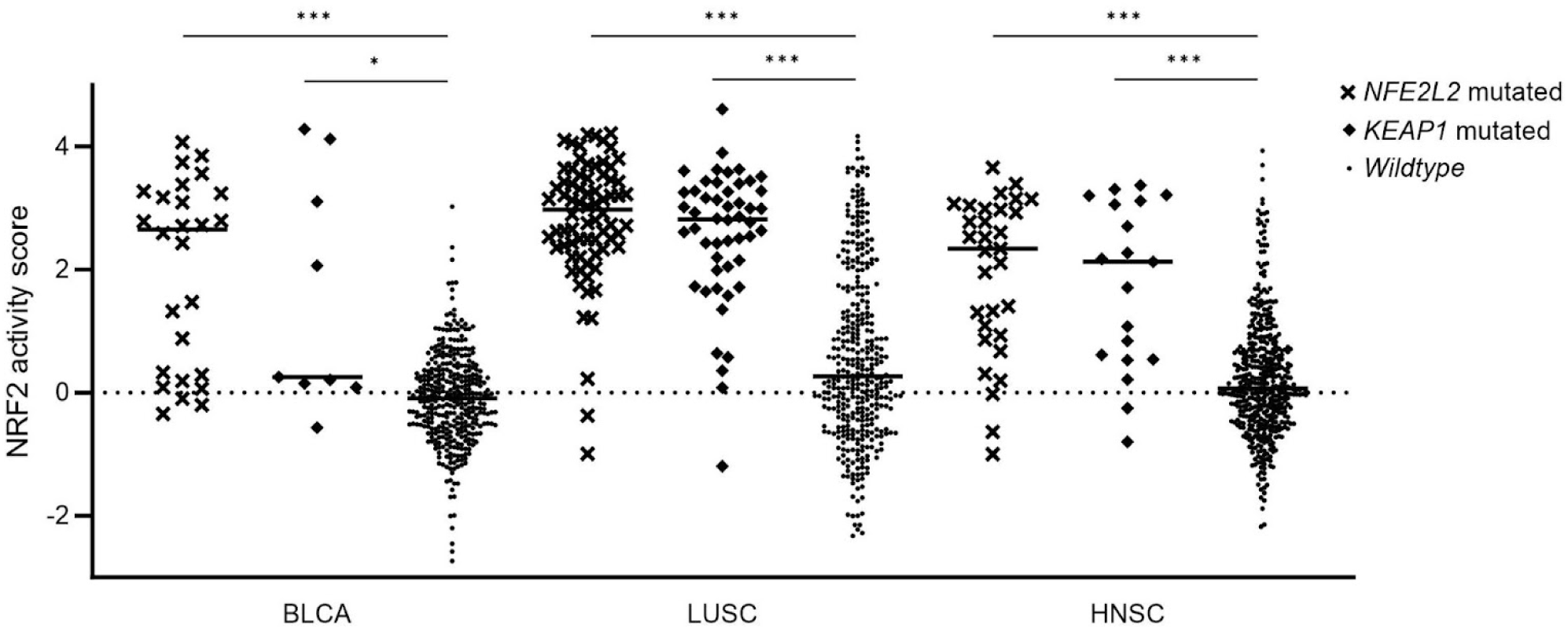
NRF2 pathway activity in bladder, lung and head & neck cancers. NRF2 pathway overactivity (as calculated in ^10^) is significantly elevated in *NFE2L2* and *KEAP1* mutated tumours (TCGA bladder carcinoma, lung squamous carcinoma, head & neck squamous carcinoma) relative to tumours without non-synonymous short nucleotide variants in these genes. However, some equally high NRF2 pathway activation is observed without mutation. Pairwise Kruskal-Wallis test significance indicated, where * p<0.05, *** p<0.001.

## References

[1] Martincorena I, Raine KM, Gerstung M, Dawson KJ, Haase K, Van Loo P, et al. Universal patterns of selection in cancer and somatic tissues. Cell. 2017;171(5):1029–1041.e21.

[2] Kamoun A, de Reyniès A, Allory Y, Sjödahl G, Robertson AG, Seiler R, et al. A consensus molecular classification of muscle-invasive bladder cancer. Eur Urol. 2020;77(4):420–433.

[3] Cross WR, Eardley I, Leese HJ, Southgate J. A biomimetic tissue from cultured normal human urothelial cells: analysis of physiological function. Am J Physiol Renal Physiol. 2005;289(2):F459–F468.

[4] Care MA, Westhead DR, Tooze RM. Parsimonious Gene Correlation Network Analysis (PGCNA): a tool to define modular gene co-expression for refined molecular stratification in cancer. NPJ Syst Biol Appl. 2019;5:13.

[5] Cotillas EA, Bernardo C, Veerla S, Liedberg F, Sjödahl G, Eriksson P. A Versatile and Upgraded Version of the LundTax Classification Algorithm Applied to Independent Cohorts. J Mol Diagn. 2024;26(12):1081–1101.

[6] DeBlasi JM, Falzone A, Caldwell S, Prieto-Farigua N, Prigge JR, Schmidt EE, et al. Distinct Nrf2 Signaling Thresholds Mediate Lung Tumor Initiation and Progression. Cancer Res. 2023;83(12):1953–1967.

[7] Rojo de la Vega M, Chapman E, Zhang DD. NRF2 and the Hallmarks of Cancer. Cancer Cell. 2018;34(1):21–43.

[8] Paik PK, Fan PD, Qeriqi B, Namakydoust A, Daly B, Ahn L, et al. Targeting NFE2L2/KEAP1 mutations in advanced NSCLC with the TORC1/2 inhibitor TAK-228. J Thorac Oncol. 2023;18(4):516–526.

[9] Robertson AG, Kim J, Al-Ahmadie H, Bellmunt J, Guo G, Cherniack AD, et al. Comprehensive molecular characterization of muscle-invasive bladder cancer. Cell. 2017;171(3):540–556.e25.

[10] Härkönen J, Pölönen P, Deen AJ, Selvarajan I, Teppo HR, Dimova EY, et al. A pan-cancer analysis shows immunoevasive characteristics in NRF2 hyperactive squamous malignancies. Redox Biol. 2023;61(102644):102644.

[11] Bray N, Pimentel H, Melsted P, Pachter L. Near-optimal RNA-Seq quantification with kallisto. Nat Biotechnol. 2016;34(5):525–527.

[12] Soneson C, Love MI, Robinson MD. Differential analyses for RNA-seq: transcript-level estimates improve gene-level inferences. F1000Res. 2015;4:1521.

[13] Peixoto TP. Nonparametric Bayesian inference of the microcanonical stochastic block model. Phys Rev E. 2017;95(1-1):012317.

[14] Peixoto TP, Kirkley A. Implicit models, latent compression, intrinsic biases, and cheap lunches in community detection. Phys Rev E. 2023;108(2-1):024309.

[15] Lambert SA, Jolma A, Campitelli LF, Das PK, Yin Y, Albu M, et al. The Human Transcription Factors. Cell. 2018;172(4):650–665.

[16] Li H, Handsaker B, Wysoker A, Fennell T, Ruan J, Homer N, et al. The Sequence Alignment/Map format and SAMtools. Bioinformatics. 2009;25(16):2078–2079.

[17] Quinlan AR, Hall IM. BEDTools: a flexible suite of utilities for comparing genomic features. Bioinformatics. 2010;26(6):841–842.

[18] Pimentel H, Bray NL, Puente S, Melsted P, Pachter L. Differential analysis of RNA-seq incorporating quantification uncertainty. Nat Methods. 2017;14(7):687–690.

[19] Xiao Y, Hsiao TH, Suresh U, Chen HIH, Wu X, Wolf SE, et al. A novel significance score for gene selection and ranking. Bioinformatics. 2014;30(6):801–807.

[20] Subramanian A, Tamayo P, Mootha VK, Mukherjee S, Ebert BL, Gillette MA, et al. Gene set enrichment analysis: a knowledge-based approach for interpreting genome-wide expression profiles. Proc Natl Acad Sci U S A. 2005;102(43):15545–15550.

[21] Liberzon A, Birger C, Thorvaldsdóttir H, Ghandi M, Mesirov JP, Tamayo P. The Molecular Signatures Database (MSigDB) hallmark gene set collection. Cell Syst. 2015;1(6):417–425.

[22] Liberzon A, Subramanian A, Pinchback R, Thorvaldsdóttir H, Tamayo P, Mesirov JP. Molecular signatures database (MSigDB) 3.0. Bioinformatics. 2011;27(12):1739–1740.

[23] Milacic M, Beavers D, Conley P, Gong C, Gillespie M, Griss J, et al. The Reactome Pathway Knowledgebase 2024. Nucleic Acids Res. 2024;52(D1):D672–D678.

[24] Cerami E, Gao J, Dogrusoz U, Gross BE, Sumer SO, Aksoy BA, et al. The cBio cancer genomics portal: an open platform for exploring multidimensional cancer genomics data. Cancer Discov. 2012;2(5):401–404.

[25] Mayakonda A, Lin DC, Assenov Y, Plass C, Koeffler HP. Maftools: efficient and comprehensive analysis of somatic variants in cancer. Genome Res. 2018;28(11):1747–1756.

[26] Suehnholz SP, Nissan MH, Zhang H, Kundra R, Nandakumar S, Lu C, et al. Quantifying the Expanding Landscape of Clinical Actionability for Patients with Cancer. Cancer Discov. 2024;14(1):49–65.

[27] Marzouka NAD, Eriksson P. multiclassPairs: an R package to train multiclass pair-based classifier. Bioinformatics. 2021;37(18):3043–3044.

[28] Dobin A, Davis CA, Schlesinger F, Drenkow J, Zaleski C, Jha S, et al. STAR: ultrafast universal RNA-seq aligner. Bioinformatics. 2013;29(1):15–21.

[29] Ewels PA, Peltzer A, Fillinger S, Patel H, Alneberg J, Wilm A, et al. The nf-core framework for community-curated bioinformatics pipelines. Nat Biotechnol. 2020;38(3):276–278.

[30] Dyer SC, Austine-Orimoloye O, Azov AG, Barba M, Barnes I, Barrera-Enriquez VP, et al. Ensembl 2025. Nucleic Acids Res. 2025;53(D1):D948–D957.

[31] McKenna A, Hanna M, Banks E, Sivachenko A, Cibulskis K, Kernytsky A, et al. The Genome Analysis Toolkit: a MapReduce framework for analyzing next-generation DNA sequencing data. Genome Res. 2010;20(9):1297–1303.

[32] Cingolani P, Platts A, Wang LL, Coon M, Nguyen T, Wang L, et al. A program for annotating and predicting the effects of single nucleotide polymorphisms, SnpEff: SNPs in the genome of Drosophila melanogaster strain w1118; iso-2; iso-3. Fly (Austin). 2012;6(2):80–92.

[33] McLaren W, Gil L, Hunt SE, Riat HS, Ritchie GRS, Thormann A, et al. The Ensembl Variant Effect Predictor. Genome Biol. 2016;17(1):122.

